# Encoding and decoding analysis of music perception using intracranial EEG

**DOI:** 10.1101/2022.01.27.478085

**Authors:** Ludovic Bellier, Anaïs Llorens, Déborah Marciano, Gerwin Schalk, Peter Brunner, Robert T. Knight, Brian N. Pasley

**Affiliations:** Helen Wills Neuroscience Institute, University of California, Berkeley, Berkeley, CA, USA; Department of Neurosurgery, Washington University School of Medicine, St. Louis, MO, USA; Department of Neurology, Albany Medical College, Albany, NY, USA; National Center for Adaptive Neurotechnologies, Albany, NY, USA; Department of Psychology, University of California, Berkeley, Berkeley, CA, USA

## Abstract

Music perception engages multiple brain regions, however the neural dynamics of this core human experience remains elusive. We applied predictive models to intracranial EEG data from 29 patients listening to a Pink Floyd song. We investigated the relationship between the song spectrogram and the elicited high-frequency activity (70-150Hz), a marker of local neural activity. Encoding models characterized the spectrotemporal receptive fields (STRFs) of each electrode and decoding models estimated the population-level song representation. Both methods confirmed a crucial role of the right superior temporal gyri (STG) in music perception. A component analysis on STRF coefficients highlighted overlapping neural populations tuned to specific musical elements (vocals, lead guitar, rhythm). An ablation analysis on decoding models revealed the presence of unique musical information concentrated in the right STG and more spatially distributed in the left hemisphere. Lastly, we provided the first song reconstruction decoded from human neural activity.

## Introduction

Music is a universal experience across all ages and cultures and is a core part of our emotional, cognitive, and social lives^1^. Understanding the neural substrate supporting music perception is a central goal in auditory neuroscience, and multiple questions remain including which musical elements (e.g., melody, harmony, rhythm) are encoded in the brain and what are the neural dynamics of brain regions underlying music perception. The last decades have seen tremendous progress in understanding the neural basis of music perception^2^, with multiple studies assessing the neural correlates of isolated musical elements such as timbre^3,4^, pitch^5,6^, melody^7,8^, harmony^9,10^ and rhythm^11,12^. These studies have established that music perception relies on a broad network of subcortical and cortical regions, including primary and secondary auditory cortices, sensorimotor areas, and inferior frontal gyri^13–16^. Both hemispheres have been shown to be involved in music processing, with a relative preference for the right hemisphere^17,18^.

These studies provide a foundation for understanding music perception. However, they typically focus on isolated musical elements or specific cortical areas. Further, they rely on brain imaging methods with either limited temporal or spatial resolution^19^ (fMRI and EEG, respectively), and on standard trial-based paradigms and analytic methods. To address these limitations, we used a naturalistic auditory stimulus listening paradigm, and applied encoding and decoding analyses to intracranial electroencephalography (iEEG) data, known for its unique spatiotemporal resolution.

We used a popular rock song (*Another Brick in the Wall, Part 1*, by Pink Floyd) as our naturalistic auditory stimulus. Studies employing restricted or synthetic stimuli are useful to assess specific aspects of auditory processing but may miss brain regions involved in higher-order processing^20,21^. Due to nonlinearities in the auditory pathways, probing the brain with isolated notes elicits neural activity in the primary auditory cortex (A1), but fails to activate areas encoding higher-order musical elements such as chords (i.e., at least three notes played together), harmony (i.e., the relationship between a system of chords), or rhythm (i.e., the temporal arrangement of notes). Using a rich and complex auditory stimulus elicits a robust and distributed neural response, allowing study of the extended neural network underlying music perception.

Music research participants are often asked to actively perform a task, such as detecting a target^3,7,8^, focusing on a particular auditory object^22,23^, or expressing a perceptual judgement^6,10^. Such tasks are necessary to study key aspects of auditory cognition, such as attention, working memory or emotions. However, the dual task nature of these approaches requiring both listening and responding distracts participants from pure music listening and confounds neural processing of music with decision processes and motor activity. To address these issues, we implemented a passive listening paradigm mimicking the everyday music-listening experience. A naturalistic music listening experience provides an uninterrupted window for assessment of higher-order aspects of musical experience (e.g., sense of beat built over time, or melodic expectations^24^) optimizing our chances at observing the full network underlying the perception of musical elements.

We recorded intracranial EEG (iEEG) data directly from the cortical surface of neurosurgical patients (electrocorticography; ECoG). This unique window on cortical processing combines the temporal resolution of electrophysiological techniques, with the spatial resolution of fMRI^25^. In addition, iEEG provides direct access to High-Frequency Activity (HFA; 70-150Hz). HFA is an index of non-oscillatory neural activity, reflecting information processing linked to local single unit firing in the infragranular cortical layers and dendritic potential in supraganular layers^26^, and to the BOLD signal in fMRI^27^. Given the direct contact between electrodes and brain tissue, iEEG benefits from an excellent signal-to-noise ratio. This is especially valuable in our naturalistic approach since it provides reliable HFA at the single-trial level enabling individual subject modeling.

We employed predictive modeling tools to take advantage of the complexity of our naturalistic stimulus and the richness of iEEG data. Specifically, we used encoding models to characterize the spectrotemporal receptive fields (STRF) of each electrode and decoding models to reconstruct the song stimulus from population neural activity. Encoding models predict neural activity at one electrode from a representation of the stimulus (e.g., spectrogram, spectrotemporal modulations, onset of notes). When this representation is a spectrogram, encoding models are called spectro-temporal receptive fields (STRFs), and a plot of their trained coefficients can be interpreted as the spectrogram of the ideal auditory stimulus to elicit an increase of neural activity at the observed electrode. These models have been successfully used to evidence key properties of the neural auditory system. This technique originated with action potential data recorded in animal models in response to artificial stimuli^28^. Recent algorithmic and machine-learning developments expanded its use to human brain imaging data and naturalistic stimuli^29^. Within the last decade, STRFs have been used to quantitatively characterize the spectrotemporal tuning profile of neural populations in response to speech or music. Notably, STRFs were used to evidence rapid plasticity of the human auditory cortex in speech perception^30^, an antero-posterior parcellation of the human superior temporal gyri^31^ (STG), and a partial overlap between the neural activity underlying music imagery and music perception^32^. By considering the full complexity of the auditory stimulus, as opposed to condition-based task design that often focuses on a single contrast dimension, and by revealing the tuning patterns of neural populations, STRFs constitute a tool of choice to investigate the neural coding supporting music perception.

Decoding models predict a representation of the stimulus from the elicited neural activity, often obtained from many electrodes. Their usage has exploded in the last decade for analyzing complex datasets without sacrificing potential dimensions of interest^29^. In the music domain, most decoding models have been used in a classification approach, for example to identity a musical piece^33^ or its genre^34,35^ from the elicited neural activity, or to estimate music-related aspects beyond the stimulus level, such as musical attention^36^ or musicianship status of the listener^37^. Another application of decoding models used in the speech domain is the stimulus reconstruction approach^38,39^, where the auditory stimulus (i.e., the sound itself) is reconstructed from the elicited neural activity. Decoding performance informs on the nature of the information represented in the recorded neural activity: if a musical element can be reconstructed, this means it was represented within the set of electrodes used as input of the decoding model. We also applied an ablation analysis, a method akin to making virtual lesions on the decoding model inputs^40,41^. We removed (or ablated) sets of predictors (here, electrodes) to assess their impact on decoding accuracy. Moreover, comparing the impact of ablating different sets of electrodes provides insights on how information is uniquely or redundantly encoded between these sets.

On the applied side, stimulus reconstruction has seen recent successes for speech decoding^42–44^. Such studies have reconstructed intelligible speech from iEEG data, using nonlinear models (deep neural networks) combined with different representations of speech including speech kinematics or the movements of vocal articulators. Here we applied stimulus reconstruction in the music domain for the first time. We investigated the extent to which a song could be reconstructed from direct brain recordings, and quantified the factors impacting decoding accuracy including model type (linear vs nonlinear) and dataset dimensionality (number of electrodes, dataset duration).

The dataset we analyzed has been the focus of previous studies, although not employing encoding and decoding models^45–49^. These studies linked several musical elements, such as sound intensity or timber, to neural activity in the posterior superior temporal gyrus (STG) or sensorimotor areas. Here, we use predictive modeling tools on iEEG data recorded from 2,668 electrodes across 29 neurological patients, who passively listened to a Pink Floyd song. We used encoding models to identify responsive cortical areas and analyze their tuning patterns and decoding models both to analyze information processing through an ablation analysis and to reconstruct the song from the elicited neural activity.

## Results

### Distribution of song-responsive electrodes

To identify electrodes encoding acoustical information about the song, we fitted STRFs for all 2,379 artifact-free electrodes in the dataset, assessing how well the HFA recorded at these sites could be linearly predicted from the song’s auditory spectrogram (Fig. 1). From a dense, bilateral, predominantly frontotemporal coverage (Fig. 2A), we identified 347 electrodes with a significant STRF (Fig. 2B). We found a higher proportion of song-responsive electrodes in the right hemisphere. There were 199 significant electrodes out of 1,479 total in the left hemisphere and 148 out of 900 in the right one (Fig. 2B, 13.5% against 16.4%, respectively; *X*^2^ (1, N=2,379) = 4.01, p = .045).

**Fig 1.**
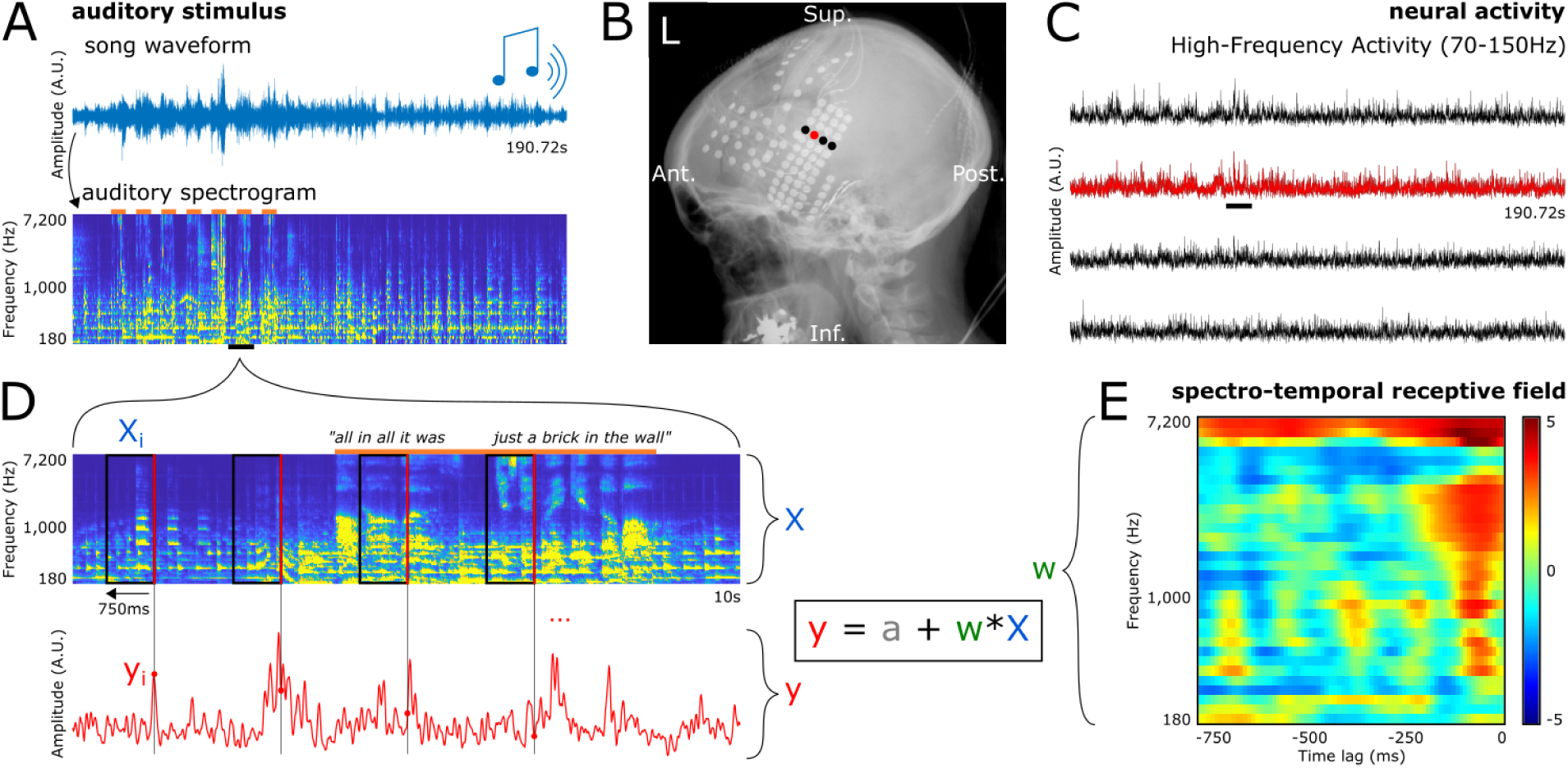
Protocol, data preparation and encoding model fitting. **A**. Top. Waveform of the entire song stimulus. Participants listened to a 190.72-second rock song (*Another Brick in the Wall, Part 1*, by Pink Floyd) using headphones. Bottom. Auditory spectrogram of the song. Orange lines on top represent parts of the song with vocals. **B**. X-ray showing electrode coverage of one representative patient. Each dot is an electrode, and the signal from the four highlighted electrodes is shown in C. **C**. HFA elicited by the song stimulus in four representative electrodes. **D**. Zoom-in on 10 seconds (black lines in A and C) of the auditory spectrogram and the elicited neural activity in a representative electrode. Each time point of the HFA (y_i_, red dot) is paired with a preceding 750-ms window of the song spectrogram (X_i_, black rectangle) ending at this time point (right edge of the rectangle, in red). The set of all pairs (X_i_, y_i_), with i ranging from .75 to 190.72 seconds, constitute the examples (or observations) used to train and evaluate the linear encoding models. Linear encoding models used here consist in predicting the neural activity (y) from the auditory spectrogram (X), by finding the optimal intercept (a) and coefficients (w). **E**. Spectro-Temporal Receptive Field (STRF) for the electrode shown in red in B, C and D. STRF coefficients are z-valued, and are represented as w in the previous equation. Note that 0 ms (timing of the observed HFA) is at the right end of the x axis, as we predict HFA from the preceding auditory stimulus.

**Fig. 2.**
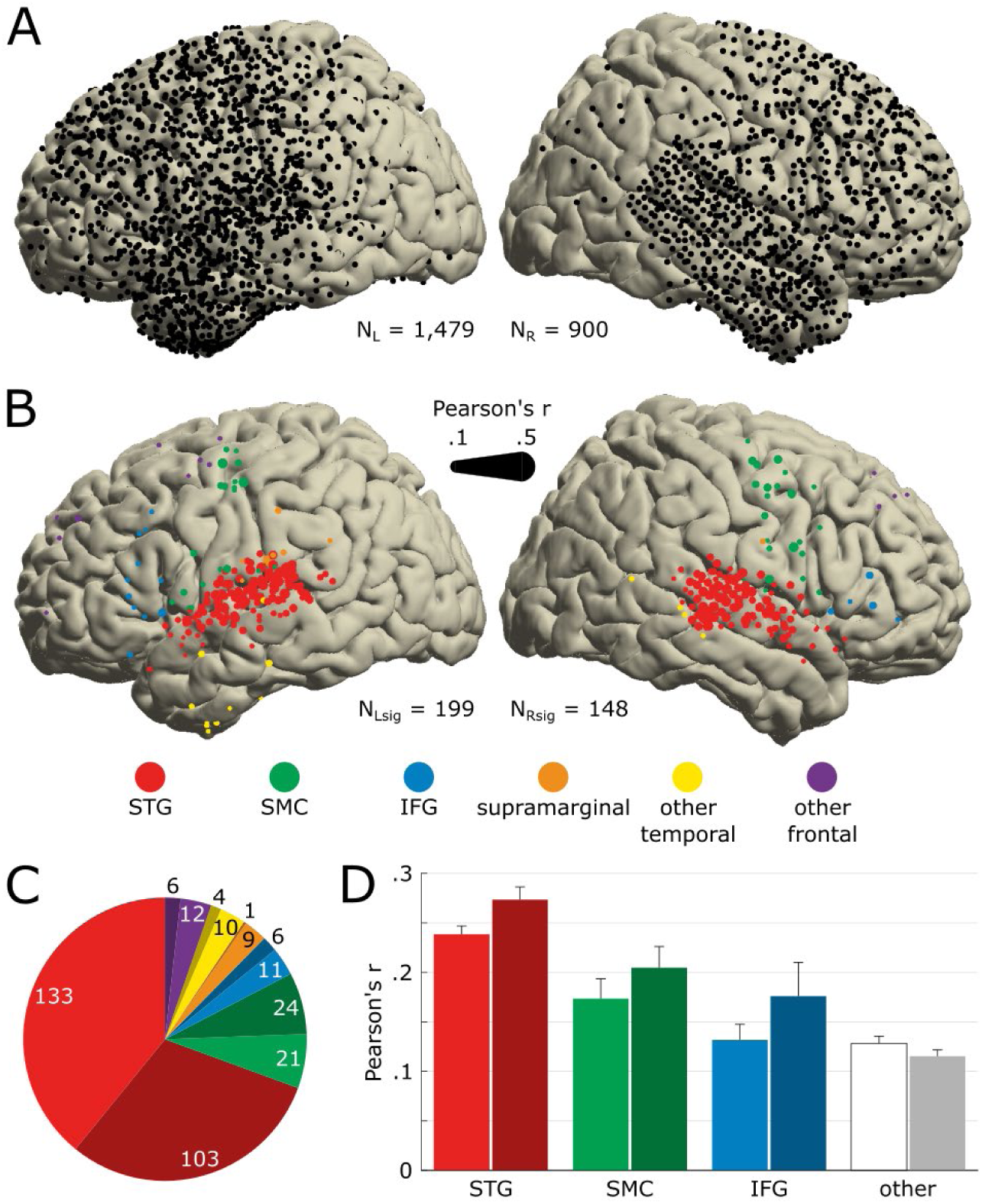
Anatomical location of song-responsive electrodes. **A**. Electrode coverage across all 29 patients shown on the MNI template (N=2,379). All presented electrodes are free of any artifactual or epileptic activity. Left hemisphere is plotted on the left. **B**. Location of electrodes significantly encoding the song’s acoustics (N_sig_=347). Significance was determined by the STRF prediction accuracy bootstrapped over 250 resamples. Marker color indicates the anatomical label as determined using the Freesurfer atlas, and marker size indicates the STRF’s prediction accuracy (Pearson’s r between actual and predicted HFA). We use the same color code in following panels and figures. **C**. Number of significant electrodes per anatomical region. Darker hue indicates a right-hemisphere location. **D**. Average STRF prediction accuracy per anatomical region. Electrodes previously labelled as *supramarginal, other temporal* (i.e., other than STG) and *other frontal* (i.e., other than SMC or IFG) are pooled together, labelled as *other* and represented in white/gray. Error bars indicate SEM.

The majority of the 347 significant electrodes (87%) were concentrated in three regions: 68% in bilateral superior temporal gyri (STG), 14.4% in bilateral sensori-motor cortices (SMC, on the pre- and postcentral gyri), and 4.6% in bilateral inferior frontal gyri (IFG; Fig. 2C). The proportion of song-responsive electrodes per region was 55.7% for STG (236 out of 424 electrodes), 11.6% for SMC (45/389), and 7.4% for IFG (17/229). The remaining 13% of significant electrodes were distributed in the supramarginal gyri and other frontal and temporal regions.

Analysis of STRF prediction accuracies (Pearson’s r) found a main effect of laterality (two-way ANOVA; *F*(1, 346) = 7.48, p = 0.0065; Fig. 2D), with higher correlation coefficients in the right hemisphere than in the left (M_R_ = .203, SD_R_ = .012; M_L_ = .17, SD_L_ = .01). We also found a main effect of cortical regions (F(3, 346) = 25.09, p < .001), with the highest prediction accuracies in STG (Tukey-Kramer post-hoc; M_STG_ = .266, SD_STG_ = .007; M_SMC_ = .194, SD_SMC_ = .017, p_STGvsSMC_ < .001; M_IFG_ = .154, SD_IFG_ = .027, p_STGvsSMC_ < .001; M_other_ = .131, SD_other_ = .016, p_STGvsSMC_ < .001). In addition, we found higher prediction accuracies in SMC compared to the group not including STG and IFG (M_SMC_ = .194, SD_SMC_ = .017; M_other_ = .131, SD_other_ = .016, p_SMCvsOther_ = .035).

### Encoding of musical elements

We analyzed STRF coefficients for all 347 significant electrodes to understand how different musical elements were encoded in different brain regions. This revealed a variety of spectrotemporal tuning patterns (Fig. 3A). To fully characterize the relationship between the song spectrogram and the neural activity, we performed an independent component analysis (ICA) on all significant STRFs. We identified three components with distinct spectrotemporal tuning patterns, each explaining more than 5% variance (Fig. 3B).

**Fig. 3.**
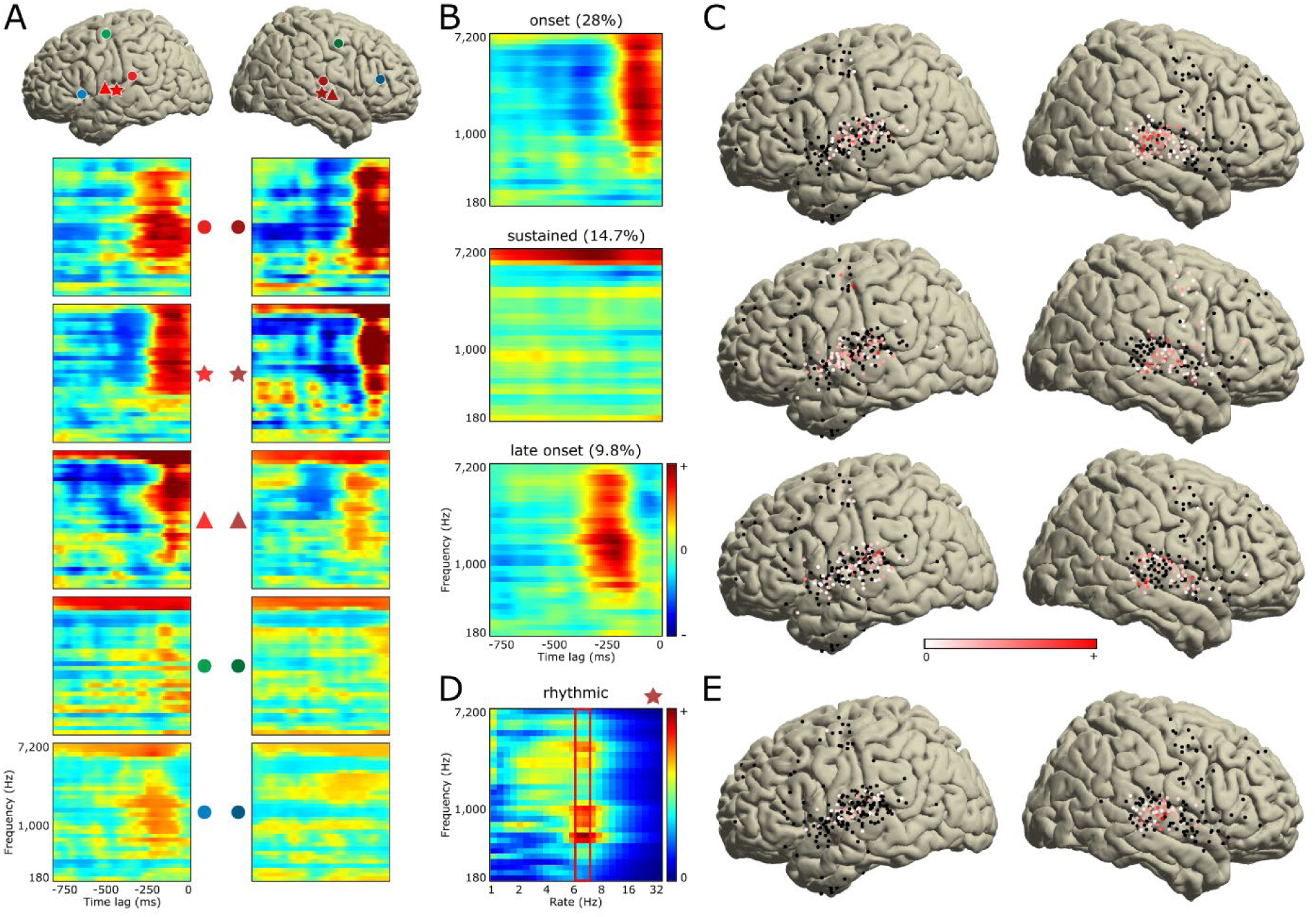
Analysis of the STRF tuning patterns. **A**. Representative set of 10 STRFs (out of the 347 significant ones) with their respective locations on the MNI template using matching markers. Color code is identical to the one used in Fig. 1. **B**. Three ICA components explaining more than 5% variance of all 347 significant STRFs. These three components show *onset, sustained* and *late onset* activity. Percentages indicate explained variance. **C**. ICA coefficients of these three components, plotted on the MNI template. Color code indicates coefficient amplitude, with STRFs of electrodes in red representing most the components. **D**. To capture tuning to the rhythm guitar pattern (16^th^ notes at 100 bpm, i.e., 6.66 Hz), pervasive throughout the song, we computed temporal modulation spectra of all significant STRFs. Example modulation spectrum is shown for a right STG electrode. For each electrode, we extracted the maximum temporal modulation value across all spectral frequencies around a rate of 6.66 Hz (red rectangle). **E**. All extracted values are represented on the MNI template. Electrodes in red show tuning to the rhythm guitar pattern.

The first component (28% explained variance) showed a cluster of positive coefficients (in red, in Fig. 3B, top row) spreading over a broad frequency range from about 500 Hz to 7 kHz, and over a narrow time window centered around 90 ms before the observed HFA (located at time lag = 0 ms, at the right edge of all STRFs). This temporally transient cluster revealed tuning to sound onsets. This component, referred to as the “onset component,” was found exclusively in electrodes located in bilateral posterior STG (Fig. 3C, top row, electrodes depicted in red). Fig. 4C, top row showed in red the parts of the song eliciting the highest HFA increase in electrodes possessing this onset component. These parts corresponded to onsets of lead guitar or synthesizer motifs (Fig. 4A, blue and purple lines, respectively; see Fig. 4E for a zoom-in) played every two bars (green lines), and to onsets of syllable nuclei in the vocals (orange lines; see Fig. 4D for a zoom-in).

**Fig. 4.**
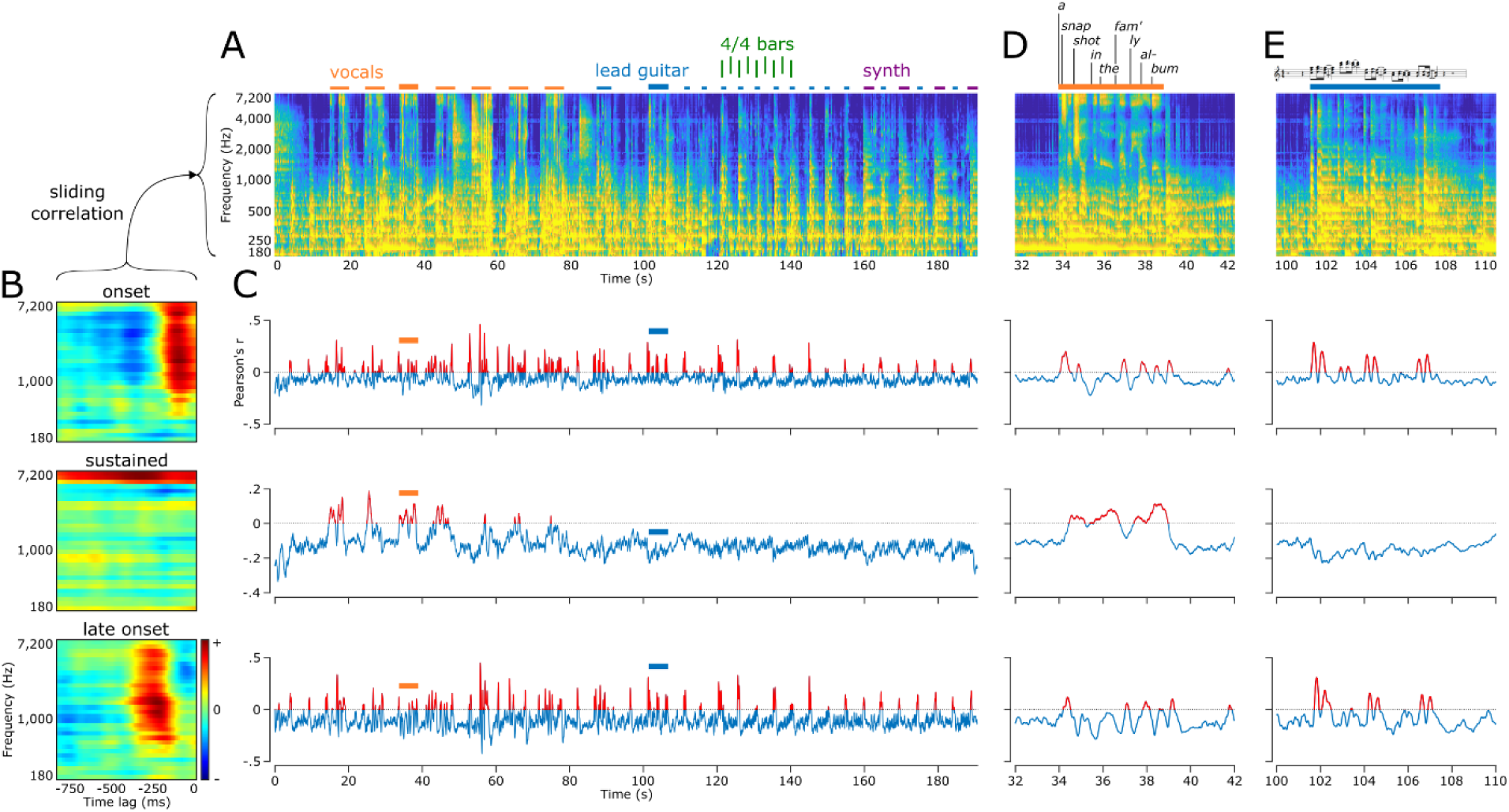
Encoding of musical elements. **A**. Auditory spectrogram of the whole song. Orange lines above the spectrogram mark all parts with vocals. Blue lines mark lead guitar motifs, and purple lines mark synthesizer motifs. Green vertical lines delineate a series of eight 4/4 bars. Thicker orange and blue lines mark locations of the zoom-ins presented in D and E, respectively. **B**. Three STRF components as presented in Fig. 3B, namely onset (top), sustained (middle) and late onset (bottom). **C**. Output of the sliding correlation between the song spectrogram (A) and each of the three STRF components (B). Positive Pearson’s r values are plotted in red, marking parts of the song that elicited an increase of HFA in electrodes exhibiting the given component. Note that for the sustained plot (middle), positive correlation coefficients are specifically observed during vocals. Also, note for both the onset and late onset plots (top and bottom, respectively), positive r values in the second half of the song corresponds to lead guitar and synthesizer motifs, occurring every other 4/4 bar. **D**. Zoom-in on the third vocals. Lyrics are presented above the spectrogram, decomposed into syllables. Most syllables triggered an HFA increase in both onset and late onset plots (top and bottom, respectively), while a sustained increase of HFA was observed during the entire vocals (middle). **E**. Zoom-in on a lead guitar motif. Sheet music is presented above the spectrogram. Most notes triggered an HFA increase in both onset and late onset plots(top and bottom, respectively), while there was no HFA increase for the sustained component (middle).

The second component (14.7% explained variance) showed a cluster of positive coefficients (in red, in Fig. 3B, middle row) spreading over the entire 750ms time window, and over a narrow frequency range from about 4.8 to 7 kHz. This component, referred to as the “sustained component,” was found in electrodes located in bilateral mid- and anterior STG, and in bilateral SMC (Fig. 3C, middle row). It correlated best with parts of the song containing vocals, thus suggesting tuning to speech (Fig. 4C, middle row, in red; see Fig. 4D for a zoom-in).

The third component (9.8% explained variance) showed a similar tuning pattern as the onset component, only with a longer latency of about 210 ms before the observed HFA (Fig. 3B, bottom row). This component, referred from now on as the “late onset component,” was found in bilateral posterior and anterior STG, neighboring the electrodes representing the onset component, and in bilateral SMC (Fig. 3C, bottom row). As with the onset component, this late onset component was most correlated with onsets of lead guitar and synthesizer motifs and of syllable nuclei in the vocals, only with a longer latency (Fig. 4C, bottom row; see Fig. 4D and 4E for zoom-ins).

A fourth component was found by computing the temporal modulations and extracting the maximum coefficient around a rate of 6.66 Hz for all 347 STRFs (Fig. 3D, red rectangle). This rate corresponded to the 16th notes of the rhythm guitar, pervasive throughout the song, at the song tempo of 99 bpm (beats per minute). It was translated in the STRFs as small clusters of positive coefficients spaced by 150 ms (1 / 6.66 Hz) from each other (e.g., Fig. 3A, electrode 5). This component, referred from now on as the “rhythmic component,” was found in electrodes located in bilateral mid STG (Fig. 3E).

### Anatomo-functional distribution of the song’s acoustic information

To assess the role of these different cortical regions and functional components in representing musical features, we performed an ablation analysis using linear decoding models. We first computed linear decoding models for each of the 32 frequency bins of the song spectrogram, using the HFA of all 347 significant electrodes as predictors. This yielded an average prediction accuracy of .62 (Pearson’s r; min .27 - max .81). We then removed (or *ablated*) anatomically- or functionally defined sets of electrodes and computed a new series of decoding models, to assess how each ablation would impact the decoding accuracy. We used prediction accuracies of the full, 347-electrode models as baseline values (Fig. 5). We found a significant main effect of electrode sets (one-way ANOVA; *F*(1, 24) = 78.4, p < .001). We then ran a series of post-hoc analyses to examine the impact of each set on prediction accuracy.

**Fig. 5.**
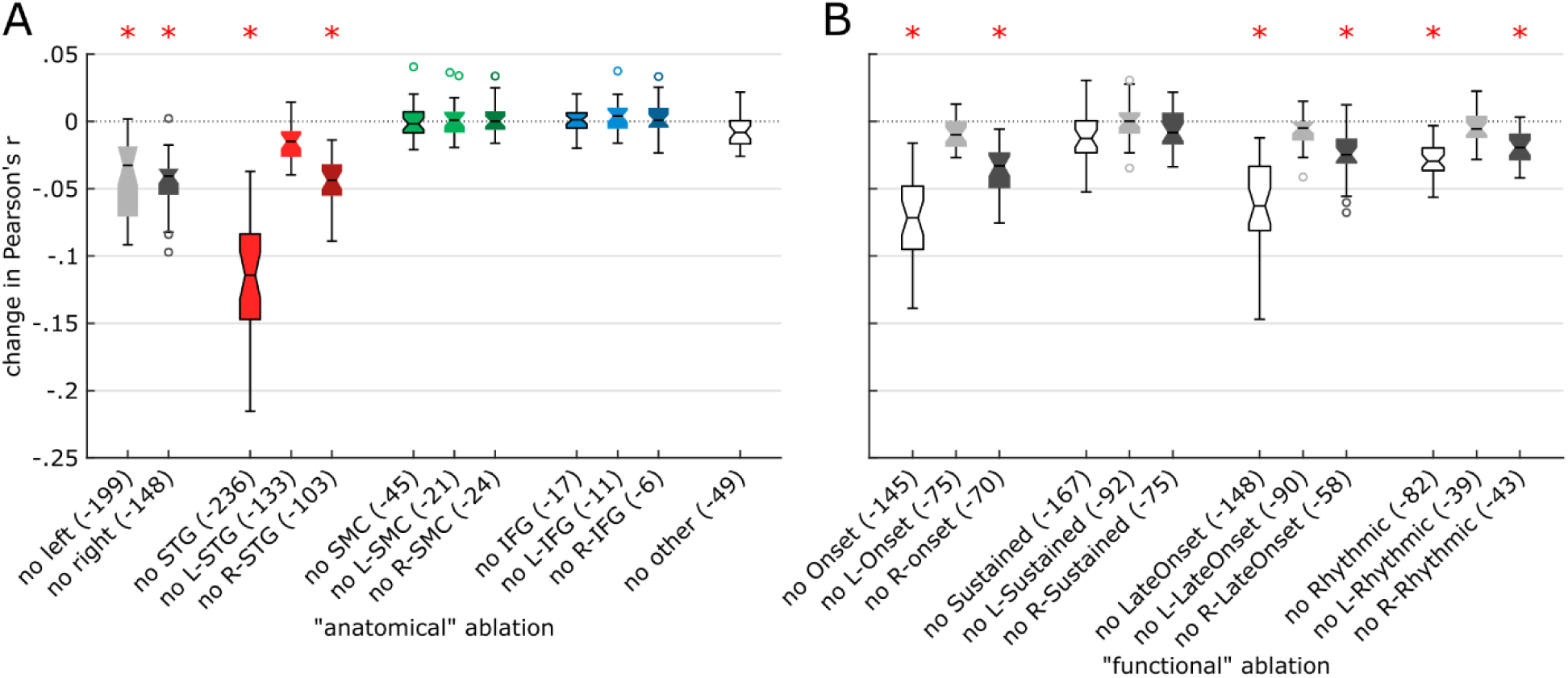
Ablation analysis on linear decoding models. We performed “virtual lesions” in the predictors of decoding models, by ablating either anatomical (**A**) or functional (**B**) sets of electrodes. Ablated sets are shown on the x axis, and their impacts on the prediction accuracy (Pearson’s r) of linear decoding models, as compared to the performance of a baseline decoding model using all 347 significant electrodes, are shown on the y axis. For each ablation, a notched box plot represents the distribution of the changes in decoding accuracy for all 32 decoding models (one model per frequency bin of the auditory spectrogram). Red asterisks indicate significant impact from ablating a given set of electrodes.

#### Anatomical ablations (Fig. 5A)

Removing all STG or all right STG electrodes impacted prediction accuracy (p < .001), with removal of all STG electrodes having the highest impact compared to all other electrode sets (p < .001). Removal of right STG electrodes had higher impact than left STG removal (p < .001), and no impact of removing left STG electrodes was found (p = .156). Together, this suggests that: 1) bilateral STG represented unique musical information compared to other regions, 2) right STG had unique information compared to left STG, and 3) part of the musical information in left STG was redundantly encoded in right STG. Ablating SMC, IFG or all other regions did not impact prediction accuracy (p > .998). Removing either all left or all right electrodes significantly reduced the prediction accuracy (p < .001), with no significant difference between all left and all right ablations (p = 1). These results suggest that both hemispheres represent unique information and contribute to song decoding. Furthermore, the fact that removing single regions in the left hemisphere had no impact but removing all left electrodes did, suggests redundancy within the left hemisphere, with musical information being spatially distributed across left hemisphere regions.

#### Functional ablations (Fig. 5B)

Removing all onset electrodes and right onset electrodes both impacted prediction accuracy (p < .001), with a highest impact for all onset (p < .001). No impact of removing left onset electrodes was found (p = .994). This suggests that right onset electrodes had unique information compared to left onset electrodes, and that part of the musical information in left onset electrodes was redundantly encoded in right onset electrodes. A similar pattern of higher right hemisphere involvement was observed with the late onset component (p < .001). Removing all rhythmic and right rhythmic electrodes both significantly impacted the decoding accuracy (p < .001 and p = .007, respectively), while we found no impact of removing left rhythmic electrodes (p = 1). We found no difference between removing all rhythmic and right rhythmic electrodes (p = .973). This suggests that right rhythmic electrodes had unique information, none of which was redundantly encoded in left rhythmic electrodes. Despite the substantial number of sustained electrodes, no impact of removing any set was found (p > .745). Note that as opposed to anatomical sets, functional sets of electrodes partially overlapped. This impeded our ability to reach conclusions regarding the uniqueness or redundancy of information *between* functional sets.

### Song reconstruction and methodological factors impacting decoding accuracy

Finally, we tested if we could reconstruct the song from neural activity, and how methodological factors such as the number of electrodes included in the model, the dataset duration or the model type at use impacted decoding accuracy. A bootstrap analysis revealed a logarithmic relationship between how many electrodes were used as predictors in the decoding model and the resulting prediction accuracy (Fig. 6A). For example, 80% of the best prediction accuracy (using all 347 significant electrodes) was obtained with 43 (or 12.4%) electrodes. A similar relationship was observed between dataset duration and prediction accuracy (Fig. 6B). For example, 90% of the best performance (using the whole 190.72s song) was obtained using 69 seconds (or 36.1%) of data.

**Fig. 6.**
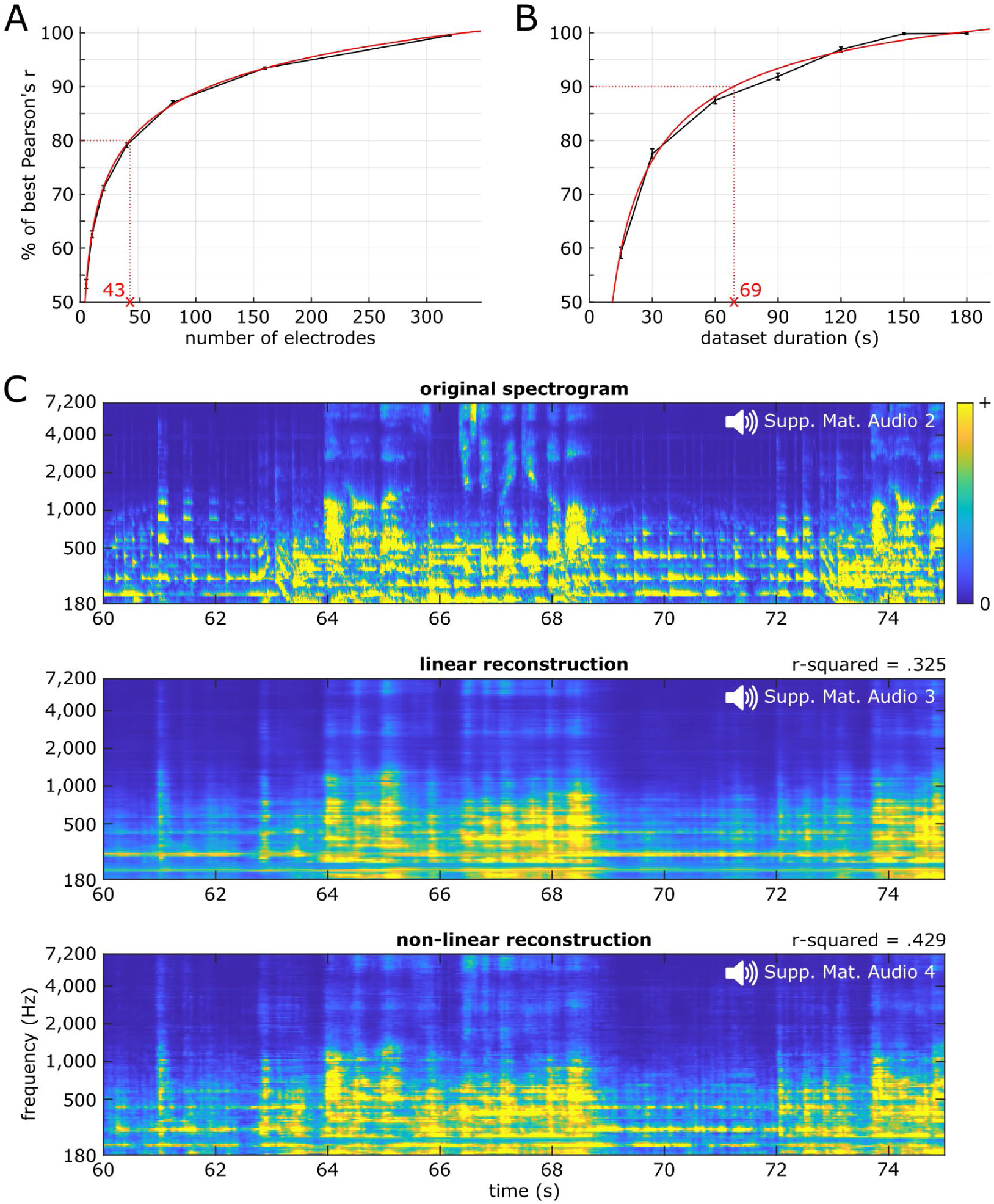
Song reconstruction and methodological considerations. **A**. Prediction accuracy as a function of the number of electrodes included as predictors in the linear decoding model. On the y axis, 100% represents the maximum decoding accuracy, obtained using all 347 significant electrodes. The black curve shows data points obtained from a 100-resample bootstrapping analysis, while the red curve shows a two-term power series fit line. **B**. Prediction accuracy as a function of dataset duration. **C**. Auditory spectrograms of the original song (top), and of the reconstructed song using either linear (middle) or nonlinear models (bottom). This 15-second song excerpt was held out during hyperparameter tuning through cross-validation and model fitting, and solely used as a test set to evaluate model performance. Corresponding audio waveforms were obtained through an iterative phase-estimation algorithm, and can be listened to in Supp. Mat. Audio 2, 3 and 4, respectively. Average effective r-squared across all 128 frequency bins is shown above both decoded spectrograms.

Regarding model type, linear decoding provided an average decoding accuracy of .325 (median of the 128 models’ effective r-squared; IQR .232), while nonlinear decoding using a two-layer, fully connected neural network (multilayer perceptron; MLP) yielded an average decoding accuracy of .429 (IQR .222). This 32% increase in effective r-squared (+.104 from .325) was significant (paired t-test, t(127) = 17.48, p < .001). In line with this higher effective r-squared for MLPs, the decoded spectrograms revealed differences between model types, with the nonlinear reconstruction (Fig. 6C, bottom row) showing finer spectro-temporal details, relatively to the linear reconstruction (Fig. 6C, middle row). Overall, the linear reconstruction (Supplementary Material Audio 3) sounded muffled with strong rhythmic cues on the presence of foreground elements (vocals syllables and lead guitar notes); a sense of spectral structure underlying timbre and pitch of lead guitar and vocals; a sense of harmony (chord progression moving from Dm to F, C and Dm); but limited sense of the rhythm guitar pattern. The nonlinear reconstruction (Supp. Mat. Audio 4) provided a recognizable song, with richer details as compared to the linear reconstruction. Especially, perceptual quality of spectral elements such as pitch and timbre were improved, and phoneme identity was perceptible. There was also a stronger sense of harmony and an emergence of the rhythm guitar pattern.

## Discussion

We applied predictive modeling analyses on iEEG data obtained from patients listening to a Pink Floyd song. Encoding models documented a central role of bilateral STG and a right-hemisphere preference in music perception. Our results revealed partially overlapping cortical areas that encoded different musical elements. An ablation analysis on decoding models showed that both the left and right hemispheres contained unique musical information, and that part of the information between left and right STG was redundant. Moreover, in the left hemisphere, we observed that musical information was spatially distributed between regions, beyond STG. On a methodological side, we quantified the impact of the number of electrodes, dataset duration and model type (linear vs nonlinear) on decoding accuracy. Notably, we provide the first recognizable song reconstruction directly decoded from human intracranial EEG data.

We observed a right hemispheric preference for music perception, with a higher proportion of electrodes with significant STRFs, higher STRF prediction accuracies, and a higher impact of ablating right electrode sets (both anatomical and functional) from the decoding models. While there was a statistical preference for the right hemisphere, left hemisphere electrodes also exhibited significant STRFs and a reduced prediction accuracy when ablated. These results are in accord with prior research, showing that music perception relies on a bilateral network, with a relative right lateralization^17,18,50^.

We also found that the spatial distribution of musical information differed between hemispheres, as suggested by the ablation results. Redundant musical information was distributed between STG, SMC and IFG in the left hemisphere, whereas unique musical information was concentrated in STG in the right hemisphere. Such spatial distribution is reminiscent of the dual-stream model of speech processing^51^. However, the absence of right SMC or IFG involvement in the ablation analysis was surprising given their reported role in music processing^16,52^. Still, we observed significant STRFs in bilateral SMC and IFG, with possible roles in encoding vocals-related information and speech or melodic syntaxis, respectively^43,53,54^.

We found a critical role of bilateral STG in representing musical information, in line with prior human studies^32,35,52,55^. As observed in other studies, STRFs obtained from the STG had rich, complex tuning patterns. To assess the anatomo-functional organization of music perception in STG, we employed a component analysis on all STRFs, which revealed four components: onset, sustained, late onset and rhythmic. The onset and sustained components were similar to those observed for speech in prior work^31,56^. Specifically, the onset component was tuned to high temporal/low spectral modulations while the sustained component was tuned to low temporal/high spectral modulations.

The onset component was tuned to a broad range of frequencies but to a narrow time window peaking at 90 ms. This latency is similar to the lag at which HFA tracked music intensity profile in Ding et al.^18^. We found that the onset component was activated by both vocals (that is, syllables) and instrumental onsets (or notes). This confirms that the onset component is not speech specific, consistent with prior work^56^ showing that reversed and spectrally rotated speech also elicited onset responses.

In contrast to the onset component, we found that the sustained component (tuned to a narrow high-frequency band but observed in a wide time window) was only activated by vocals. As seen in prior work^31,56^ we observed these two components in anatomically distinct STG subregions, with the onset component in posterior STG and the sustained component in mid- and anterior STG. Interestingly, we observed single electrodes representing both the onset and the sustained components, which were mostly located in mid STG. This was not found in previous studies, likely due to the use of different data-driven approaches (clustering vs ICA). Surprisingly, in our functional ablation analysis, removing all electrodes representing the sustained component did not impact decoding accuracy, despite their substantial number (167 out of 347). This might be due to the fact that as the song is dominated by instrumentals, removing a component related to vocals had negligible impact on the decoding accuracy.

In addition to the onset and sustained component, we found evidence for two other distinct components: late onset and rhythmic. The late onset component was found in electrodes neighboring the onset component in STG and had similar tuning properties as the onset component, only peaking at a later latency of 210ms. This is in line with the findings of Nourski et al.^57^, who, using click trains and a speech syllable, observed a concentric spatial gradient of HFA onset latencies in STG, with shorter latencies in post-/mid-STG and longer latencies in surrounding tissue. Further studies are needed to understand better the relationship between the onset and late onset components, as their similar functional behavior despite such different latencies appears as a discrepancy. The rhythmic component, tuned to the 6.66 Hz sixteenth notes of the rhythm guitar, was observed in mid STG, especially in electrodes representing both onset and sustained components. This provides a novel link between HFA and a specific rhythmic signature in a subregion of STG, and extends prior studies that found an involvement of STG in a range of rhythmic processes, i.e., beat perception^58^, omissions^59^, periodicity^60^. Altogether, these four components paint a rich picture of the anatomo-functional organization of complex sound processing in the human STG.

On the methodological side, we observed a logarithmic relationship between decoding accuracy and the number of electrodes (a proxy for electrode density) or dataset duration, in line with previous literature for speech stimuli^39,42^. We showed that 80% of the maximum observed decoding accuracy was achieved with 43 electrodes or in 37 seconds, which supports the feasibility of using predictive modeling approaches in relatively small datasets. Interestingly, ablating the 167 sustained electrodes (Fig. 5B) had no significant impact on decoding accuracy, while ablating the 43 right rhythmic electrodes did. This observation shows that electrode functional role and anatomical location were primordial factors.

We reconstructed a recognizable song using nonlinear models predicting the song’s acoustics from the elicited HFA. Linear decoding provided a surprisingly good r-squared of 32.5% explained variance but nonlinear reconstruction performed better at all levels, with a higher r-squared of 42.9%, a more detailed decoded spectrogram, and a recognizable song. This is likely due to the multilayer perceptron’s ability to decode nonlinearly transformed acoustic information represented in non-primary auditory areas such as STG^61^. Decoding the song spectrogram from electrodes in primary auditory cortices (A1, accessible with stereotactic EEG/depth electrodes) might improve the performance of linear models. While nonlinear reconstruction performed better than linear reconstruction, it lacked clarity on some musical elements, especially on the background rhythm guitar pattern. This might be due to several limiting factors: dataset duration could be too short (only slightly more than three minutes) to fully train MLPs; musical information represented in STG could be too nonlinearly transformed, with information loss irreversible even using MLPs; the rhythm guitar pattern, pervasive throughout the song and played in the background, might be perceived as less relevant than vocals or lead guitar phrases, leading to less representation in higher-order auditory areas; lastly, being of lower amplitude than vocals or lead guitar notes in the spectrogram, the rhythm guitar could contribute less to the Mean Squared Error during model fitting, leading to reduced reconstruction.

An important open question is whether there exist brain regions and networks that are specific to music, or whether music-related information is processed in input agnostic auditory pathways^50,62,63^. While this study links musical elements to STRF components and precise anatomical locations, it is unlikely that these regions respond specifically to music. Rather our findings suggest non-music-specific encoding of musical elements. The fact that onset and late onset components responded to syllables, lead guitar and synthesizer (Fig. 4) suggests that subparts of STG process both vocals and music. Although one could argue that the rhythmic component (Fig. 3D and E) is music specific as it is clearly related to the 6.66 Hz sixteenth notes of the rhythm guitar, this same rhythmic component shows diffuse energy between 2 and 8 Hz in the temporal modulation spectrum (Fig. 3D), compatible with syllabic rhythm^64^. On the other hand, a specificity for speech is suggested by the sustained component, as it is only activated by vocals (Fig. 4C, D and E).

Our study had several limitations. Importantly, the encoding models we used in this study to investigate the neural dynamics of music perception estimated the linear relationship between song’s acoustics and elicited HFA. It is possible that regions not highlighted by our study respond to the song, either in other neural frequency bands, or encoding higher-order musical information. Another limitation was the short duration of the song, and its limited spectrotemporal variability. More data would enhance statistical power and enable the use of more complex nonlinear models. Finally, we lacked patient-related information about musicianship status or degree of familiarity with the song, preventing us to investigate inter-individual variability.

Combining a naturalistic paradigm, unique iEEG data and novel modeling-based analyses, this study extends our knowledge of the neural dynamics underlying music perception at two levels. At the brain level, we observed a right-hemisphere preference and a preponderant role of bilateral STG in representing the song’s acoustics. Within bilateral STG, we observed partially overlapping neural populations tuned to distinct musical elements. An ablation analysis revealed the presence of unique musical information in both hemispheres, spatially distributed in the left hemisphere between STG, SMC and IFG, and concentrated in STG in the right hemisphere. At a methodological level, we showed the feasibility of applying predictive modeling on a relatively short dataset and quantified the impact of different methodological factors on the prediction accuracy of decoding models. To our knowledge, we provide the first recognizable song reconstructed from direct brain recordings. Future studies could investigate different representations of the song (i.e., notes, chords, sheet music) and different neural frequency bands (e.g., theta, alpha, beta power), and will add another brick in the wall of our understanding of music processing in the human brain.

## Methods

### Participants

Twenty-nine patients with pharmacoresistant epilepsy participated in the study. All had intracranial grids or strips of electrodes (electrocorticography, ECoG) surgically implanted to localize their epileptic foci, and electrode location was solely guided by clinical concern. Recordings took place at the Albany Medical Center (Albany, NY). All patients volunteered and gave their informed consent prior to participating in the study. The experimental protocol has been approved by the Institutional Review Boards of both the Albany Medical Center and the University of California, Berkeley. All patients had self-declared normal hearing.

### Task

Patients passively listened to the song *Another Brick in the Wall, Part 1*, by Pink Floyd (released on the album The Wall, Harvest Records/Columbia Records, 1979). They were instructed to listen attentively to the music, without focusing on any special detail. Total song duration was 190.72 seconds (waveform is represented in Fig. 1A, top; listen to Supplementary Material Audio 1 for a 15-second excerpt). The auditory stimulus was digitized at 44.1 kHz and delivered through in-ear monitor headphones (bandwidth 12Hz-23.5kHz, 20dB isolation from surrounding noise) at a comfortable sound level adjusted for each patient (50 to 60 dB SL). Eight patients had more than one recording of the present task, in which cases we selected the cleanest one (i.e., containing the least epileptic activity or noisy electrodes).

### Intracranial recordings

Direct cortical recordings were obtained through grids or strips of platinum-iridium electrodes (Ad-Tech Medical, Oak Creek, WI), with center-to-center distances of 10 mm for 21 patients, 6 mm for four, 4 mm for three or 3 mm for one. We recruited patients in the study if their implantation map covered at least partially the superior temporal gyri (left or right). The cohort consists of 28 unilateral cases (18 left, 10 right) and one bilateral case. Total number of electrodes across all 29 patients was 2,668 (range 36-250, mean 92 electrodes). ECoG activity was recorded at a sampling rate of 1,200 Hz using g.USBamp biosignal acquisition devices (g.tec, Graz, Austria) and BCI2000^65^.

### Preprocessing – Auditory stimulus

To study the relationship between the acoustics of the auditory stimulus and the ECoG-recorded neural activity, the song waveform was transformed into a magnitude-only auditory spectrogram using the NSL Matlab Toolbox^66^. This transformation mimics the processing steps of early stages of the auditory pathways, from the cochlea’s spectral filter bank to the midbrain’s reduced upper limit of phase-locking ability, and outputs a psychoacoustic-, neurophysiologic-based spectrotemporal representation of the song. The resulting auditory spectrogram has 128 frequency bins from 180 to 7,246 Hz, with characteristic frequencies uniformly distributed along a logarithmic frequency axis (24 channels per octave), and a sampling rate of 100 Hz. This full-resolution, 128-frequency-bin spectrogram is used in the song reconstruction analysis. For all other analyses, to decrease the computational load and the number of features, we outputted a reduced spectrogram with 32 frequency bins from 188 to 6,745 Hz (Fig. 1A, bottom).

### Preprocessing – ECoG data

We used the High-Frequency Activity (HFA; 70 to 150 Hz) as an estimate of local neural activity^67^ (Fig. 1C). For each dataset, we visually inspected raw recorded signals and removed electrodes exhibiting noisy or epileptic activity, with the help of a neurologist (RTK). We then extracted data aligned with the song stimulus, adding 10 seconds of data padding before and after the song (to prevent filtering-induced edge artifacts). We filtered out power-line noise, using a range of notch filters centered at 60 Hz and harmonics up to 300 Hz (Butterworth, 4^th^ order, 2 Hz bandwidth), and removed slow drifts with a 1 Hz high-pass filter (Butterworth, 4^th^ order). We used a bandpass-Hilbert approach^68^ to extract HFA, with 20-Hz-wide sub-bands spanning from 70 to 150 Hz in 5 Hz steps (70 to 90, 75 to 95, … up to 130 to 150 Hz). We chose a 20 Hz bandwidth to enable the observation of temporal modulations up to 10 Hz^69^, encompassing the 6.66 Hz sixteenth-note rhythm guitar pattern, pervasive throughout the song. This constitutes a crucial methodological point, enabling the observation of the rhythmic component (Fig. 3D). For each sub-band, we first bandpass-filtered the signal (Butterworth, 4^th^ order), then performed median-based Common Average Reference (CAR; Liu et al., 2015), and computed the Hilbert transform to obtain the envelope. We standardized each sub-band envelope using robust scaling on the whole time period (subtracting the median and dividing by the interquartile range between the 10^th^ and 90^th^ percentiles), and average them together to yield the HFA estimate. We performed CAR separately for electrodes plugged on different splitter boxes to optimize denoising in 14 participants. Finally, we removed the 10-second pads, down-sampled data to 100 Hz to match the stimulus spectrogram’s sampling rate, and tagged outlier time samples exceeding seven standard deviations for later removal in the modeling preprocessing. We used Fieldtrip^71^ (version from May 11, 2021) and homemade scripts to perform all above preprocessing steps. Unless specified otherwise, all further analyses and computations were implemented in MATLAB (The MathWorks, Natick, MA, USA; version 2021a). Code is available upon request.

### Preprocessing – Anatomical data

We followed the anatomical data processing pipeline presented in Stolk et al.^72^ to localize electrodes from a pre-implantation MRI, a post-implantation CT scan and coverage information mapping electrodes to channel numbers in the functional data. After co-registration of the CT scan to the MRI, we performed brain-shift compensation with a hull obtained using scripts from the iso2mesh toolbox^73,74^. Cortical surfaces were extracted using the Freesurfer toolbox^75^. We used volume-based normalization to convert patient-space electrode coordinates into MNI coordinates for illustration purposes, and surface-based normalization using the Freesurfer’s fsaverage template to automatically obtain anatomical labels from the aparc+aseg atlas. Labels were then confirmed by a neurologist (RTK).

### Encoding – Data preparation

We used Spectro-Temporal Receptive Fields (STRFs) as encoding models, with the 32 frequency bins of the stimulus spectrogram as features or predictors, and the HFA of a given electrode as target to be predicted.

We log-transformed the auditory spectrogram to compress all acoustic features into the same order of magnitude (e.g., low-sound-level musical background and high-sound-level lyrics). This ensured modeling would not be dominated by high-volume musical elements.

We then computed the feature lag matrix from the song’s auditory spectrogram (Fig. 1D). As HFA is elicited by the song stimulus, we aim at predicting HFA from the preceding song spectrogram. We chose a time window between 750 ms and 0 ms before HFA, to allow a sufficient temporal integration of auditory-related neural responses, while ensuring a reasonable features-to-observations ratio to avoid overfitting. This resulted in 2,400 features (32 frequency bins by 75 time lags at a sampling rate of 100 Hz).

We obtained 18,898 observations per electrode, each one consisting of a set of one target HFA value and its preceding 750-ms auditory spectrogram excerpt (19,072 samples of the whole song, minus 74 samples at the beginning for which there is no preceding 750-ms window).

At each electrode, we rejected observations for which the HFA value exceeded seven standard deviations (Z units), resulting in an average rejection rate of 1.83% (min 0% - max 15.02%, SD 3.2%).

### Encoding – Model fitting

To obtain a fitted STRF for a given electrode, we iterated through the following steps 250 times.

We first split the dataset into training, validation and test sets (60-20-20 ratio, respectively) using a custom group-stratified-shuffle-split algorithm (based on the StratifiedShuffleSplit cross-validator in scikit-learn). We defined relatively long, 2-second groups of consecutive samples as indivisible blocks of data. This ensured that training and test sets would not contain neighbor, virtually identical samples (as both music and neural data are highly correlated over short periods of time), and was critical to prevent overfitting. We used stratification to enforce equal splitting ratios between the vocal (13 to 80 s) and instrumental parts of the song. This ensured stability of model performance across all 250 iterations, by avoiding that a model could be trained on the instrumentals only and tested on the vocals. We used shuffle splitting, akin to bootstrapping with replacement between iterations, which allows us to determine test set size independently from the number of iterations (as opposed to KFold cross-validation).

We then standardized the features, by fitting a robust scaler to the training set only (estimates the median and the 2-98 quantile range; RobustScaler in sklearn package), and using it to transform all training, validation and test sets. This gives comparable importance to all features, i.e., every time lag and frequency of the auditory spectrogram.

We employed linear regression with RMSProp optimizer for efficient model convergence, Huber loss cost function for robustness to outlier samples, and early stopping to further prevent overfitting. In early stopping, a generalization error is estimated on the validation set at each training step, and model fitting ends after this error stops diminishing for 10 consecutive steps. This model was implemented in Tensorflow 1.6 and Python 3.6. The learning rate hyperparameter of the RMSProp optimizer was manually tuned to ensure fast model convergence all by avoiding exploding gradients (overshooting of the optimization minimum).

We evaluated prediction accuracy of the fitted model by computing both the correlation coefficient (Pearson’s r) and the R-squared between predicted and actual test-set target (i.e., HFA at a given electrode). Along with these two performance metrics, we also saved the fitted model coefficients.

Then, we combined these 250 split-scale-fit-evaluate iterations in a bootstrap-like approach to obtain one STRF and assess its significance (i.e., whether we can linearly predict HFA, at a given electrode, from the song spectrogram). For each STRF, we z-scored each coefficient across the 250 models (Fig. 1E). For the prediction accuracy, we computed the 95% confidence interval (CI) from the 250 correlation coefficients, and deemed an electrode as significant if its 95% CI did not contain 0. As an additional criterion, we rejected significant electrodes with an average R-squared (across the 250 models) at or below 0.

### Encoding – Analysis of prediction accuracy

To assess how strongly each brain region encodes the song, we performed a two-way ANOVA on the correlation coefficients of all electrodes showing a significant STRF, with laterality (left or right hemisphere) and area (STG, sensorimotor, IFG or other) as factors. We then performed a multiple comparison (post hoc) test to disentangle any differences between factor levels.

### Encoding – Analysis of model coefficients

We analyzed the STRF tuning patterns using an independent component analysis (ICA), to highlight electrode populations tuned to distinct STRF features. Firstly, we ran an ICA with 10 components on the centered STRF coefficients, to identify components individually explaining more than 5% of variance. We computed explained variance by back-projecting each component and using the following formula: pvaf_i_ = 100 – 100*mean(var(STRF - backproj_i_))/mean(var(STRF)); with i from 1 to 10 components, pvaf_i_ being the percentage of variance accounted for by ICA component i, STRF being the centered STRF coefficients, and backproj_i_ being the back-projection of ICA component i in electrode space. We found 3 ICA components explaining more than 5% of variance. To optimize the unmixing process, we ran a new ICA asking for three components. Then, we determined each component sign by setting as positive the sign of the most salient coefficient. Lastly, for each ICA component, we defined electrodes as representing the component if their ICA coefficient was positive.

To look at rhythmic tuning patterns, we computed the temporal modulations of each STRF. Indeed, due to their varying frequencies and latencies, they were not captured by the combined component analysis. We quantified temporal modulations between 1 and 16 Hz over the 32 spectral frequency bins of each STRF, and extracted the maximum modulation value across all 32 frequency bins between 6 and 7 Hz of temporal modulations, corresponding to the song rhythmicity of 16th notes at 99 bpm. We defined electrodes as representing the component if their maximum modulation value was above a manually defined threshold of .3.

### Encoding – Musical elements

To link STRF components to musical elements in the song, we ran a sliding-window correlation between each component and the song spectrogram. Positive correlation values indicate specific parts of the song or musical elements (i.e., vocals, lead guitar…) that elicit an increase of HFA.

### Decoding - Ablation analysis

To assess the contribution of different brain regions and STRF components in representing the song, we performed an ablation analysis. We quantified the impact of ablating sets of electrodes on the prediction accuracy of a linear decoding model computed using all 347 significant electrodes. Firstly, we constituted sets of electrodes based on anatomical or functional criteria. We defined 12 anatomical sets by combining two factors – area (whole hemisphere, STG, SMC, IFG, or other areas) and laterality (bilateral, left or right). We defined 12 functional sets by combining two factors – STRF component identified in the STRF coefficient analyses (onset, sustained, late onset, and rhythmic) and laterality (bilateral, left or right). See Fig. 5 for the exact list of electrode sets. Secondly, we computed the decoding models using the same algorithm as for the encoding models. Decoding models aim at predicting the song spectrogram from the elicited neural activity. Here, we used HFA from a set of electrodes as input, and a given frequency bin of the song spectrogram as output. For each of the 24 ablated sets of electrodes, we obtained 32 models (one per spectrogram frequency bin), and compared each one of them to the corresponding baseline model computed using all 347 significant electrodes (repeated-measure one-way ANOVA). We then performed a multiple comparison (post hoc) test to assess differences between ablations.

We based our interpretation of ablations results on the following assumptions. Collectively, as they had significant STRFs, all 347 significant electrodes represent acoustic information on the song. If ablating a set of electrodes resulted in a significant impact on decoding accuracy, we considered that this set represented unique information. Indeed, were this information shared with another set of electrodes, a compensation-like mechanism could occur and void the impact on decoding accuracy. If ablating a set of electrodes resulted in no significant impact on decoding accuracy, we considered that this set represented redundant information, shared with other electrodes (as the STRFs were significant, we ruled out the possibility that it could be because this set did not represent any acoustic information). Also, comparing the impact of a given set and one of its subsets of electrodes provided further insights on the unique or redundant nature of the represented information.

### Decoding – Parametric analyses

We quantified the influence of different methodological factors (number of electrodes, dataset duration, and model type) on the prediction accuracy of decoding models. In a bootstrapping approach, we randomly constituted subsets of 5, 10, 20, 40, 80, 160 and 320 electrodes (sampling without replacement) to be used as inputs of linear decoding models. We processed 100 bootstrap resamples (i.e., 100 sets of 5 electrodes, 100 sets of 10 electrodes…), and normalized for each of the 32 frequency bins the resulting correlation coefficients by the correlation coefficients of the full, 347-electrode decoding model. For each resample, we averaged the correlation coefficients from all 32 models (1 per frequency bin of the song spectrogram). This yielded 100 prediction accuracy estimates per number of electrodes. We then fitted a two-term power series model to these estimates, to quantify the apparent power-law behavior of the obtained bootstrap curve. We adopted the same approach for dataset duration, with excerpts of 15, 30, 60, 90, 120, 150 and 180 consecutive seconds.

To investigate the impact of model type on decoding accuracy and to assess the extent to which we could reconstruct a recognizable song, we trained linear and nonlinear models to decode each of the 128 frequency bins of the full spectral resolution song spectrogram from HFA of all 347 significant electrodes. We used the multilayer perceptron (MLP)—a simple, fully connected neural network, as nonlinear model (MLPRegressor in sklearn). We chose a MLP architecture of two hidden layers of 64 units each, based both on an extension of the *Universal Approximation Theorem* stating that a two hidden layer MLP can approximate any continuous multivariate function^76^ and on a previous study with a similar use case^42^. Since MLP layers are fully connected (i.e., each unit of a layer is connected to all units of the next layer), the number of coefficients to be fitted is drastically increased relatively to linear models (in this case, F*N + N*N + N vs F, respectively, where the total number of features F = E*L, with E representing the number of significant electrodes included as inputs of the decoding model, and L the number of time lags; and N represents the number of units per layer). Given the limited dataset duration, we reduced time lags to 500ms based on the absence of significant activity beyond this point in the STRF components, and used this L value in both linear and nonlinear models.

We defined a fixed, 15-second continuous test set during which the song contained both vocals and instrumentals (Supp. Mat. Audio 1), and held it out during hyperparameter tuning and model fitting. We tuned model hyperparameters (learning rate for linear models, and L2-regularization alpha for MLPs) through 10-resample cross-validation. We performed a grid search on each resample (i.e., training/validation split), and saved for each resample the index of the hyperparameter value yielding the minimum validation mean squared error (MSE). Candidate hyperparameter values ranged between .001 and 100 for the learning rate of linear models, and between .01 and 100 for the alpha of MLPs. We then rounded the mean of the ten resulting indices to obtain the cross-validated, tuned hyperparameter. As a homogeneous presence of vocals across training, validation and test sets was crucial for proper tuning of the alpha hyperparameter of MLPs, we increased group size to 5 seconds, equivalent to about two musical bars, in the group-stratified-shuffle-split step (see Encoding models – Model Fitting for a reference), and used this value for both linear and nonlinear models. For MLPs specifically, as random initialization of coefficients could lead to convergence towards local optima, we adopted a best-of-3 strategy where we only kept the “winning” model (i.e., yielding the minimum validation MSE) amongst three models fitted on the same resample.

Once we obtained the tuned hyperparameter, we computed 100 models on distinct training/validation splits, also adopting the best-of-3 strategy for the nonlinear models (this time keeping the model yielding the maximum test r-squared). We then sorted models by increasing r-squared, and evaluated the “effective” r-squared by computing the r-squared between the test set target (the actual amplitude time course of the song’s auditory spectrogram frequency bin) and averages of n models, with n varying from 100 to 1 (i.e., effective r-squared for the average of all 100 models, for the average of the 99 best, …, of the 2 best, of the best model). Lastly, we selected n based on the value giving the best effective r-squared, and obtained a predicted target along with its effective r-squared as an estimate of decoding accuracy. The steps above were performed for all 128 frequency bins of the song spectrogram, both for linear and nonlinear models, and we compared the resulting effective r-squared using a paired t-test.

### Decoding – Song waveform reconstruction

To explore the extent to which we could reconstruct the song from neural activity, we collected the 128 predicted targets for both linear and MLP decoding models as computed above, therefore assembling the decoded auditory spectrograms. To denoise and improve sound quality, we rose all spectrogram samples to the power of two, thus highlighting prominent musical elements such as vocals or lead guitar chords, relatively to background noise. As both magnitude and phase information are required to reconstruct a waveform from a spectrogram, we used an iterative phase-estimation algorithm to transform the magnitude-only decoded auditory spectrogram into the song waveform (*aud2wav*^66^). To have a fair basis against which we could compare the song reconstruction of the linearly and nonlinearly decoded spectrograms, we transformed the original song excerpt corresponding to the fixed test set into an auditory spectrogram, discarded the phase information, and applied this algorithm to revert the spectrogram into a waveform (Supp. Mat. Audio 2). We performed 500 iterations of this aud2wav algorithm, enough to reach a plateau where error did not improve further.

## Acknowledgements

This work has been supported by the Fondation Pour l’Audition (FPA RD-2015-2, LB), the National Institutes of Health NIH/NIBIB (R01-EB026439, P41-EB018783, PB), NIH/NINDS (U24-NS109103, U01-NS108916, R13-NS118932, PB; R01-NS21135, RTK), NIH/NIDCD (1R21-DC018374, BNP), and the National Science Foundation (NSF 1850687, BNP). We thank Stephanie Martin, Christopher R. Holdgraf, Christian Mikutta, Randolph F. Helfrich and Arjen Stolk for their helpful comments on data analysis.

## Author contributions

Study design and data acquisition (GS, PB), data preprocessing and analysis (LB, BNP), writing (LB, AL, DM), editing (RTK, BP).

## Competing interests

The authors confirm that there are no relevant financial or non-financial competing interests to report.

## Reference gender statistics

Across all 76 references, five had females as first and last authors, eight had a male first author and a female last author, 17 had a female first author and a male last author, and 38 had males as first and last authors. Eight papers had a single author, amongst which one was written by a female.

## Notes

### Competing Interest Statement

The authors have declared no competing interest.

### Summary of Updates

Updated the song reconstruction part with our latest results; added supplemental audio files; provided cosmetic adjustments to figures; improved figure legends.

